# Identifying Family Structures from Obituaries and Matching them to Patients in an Electronic Heath Record

**DOI:** 10.1101/2024.11.26.625445

**Authors:** John Mayer, Brooke Delgoffe, Scott Hebbring

## Abstract

**Motivation:** Family data is a valuable data source in bioinformatic research. This is because family members often share common genetic and environmental exposures. Collecting this family data is traditionally very labor intensive but advances in electronic health record (EHR) data mining has proven useful when identifying pedigrees linked to longitudinal health histories. These are called e-pedigrees. Unfortunately, e-pedigrees tend to miss the oldest generations who inherently have the longest and richest health histories. A good source of family data from older generations includes obituaries, as they have a formulaic nature making them a good candidate for natural language processing that can extract relationships to the decedent. While there have been several studies on obtaining such data from obituaries, we demonstrate for the first-time approaches that tie that information to an EHR.

**Results:** NLP extraction resulted in 8,166,534 family members being abstracted from 567,279 obituaries published in the state of Wisconsin. After matching decedent and family members to patients in the EHR, we identified 109,365 unique patients that were put in 34,158 pedigrees. The largest pedigree consisted of 21 individuals. Heritability of adult height was quantified (H^2^= 0.51 +- .04, P=< 1.00e-07) demonstrating this data’s use in genetic research. The heritability data, coupled with overlapping data in a biobank, suggested 80% - 90% of familial relationships were accurately defined. The totality of these findings demonstrate obituaries with the oldest generations can be highly informative for bioinformatic research.

**Availability and Implementation:** Code is available on GitHub at https://github.com/jgmayer672/ObituaryNLP.

## Introduction

Family data can be extremely valuable in biomedical research and clinical care. This is because nearly all diseases have a familial component that is influenced by shared genetics and environment (Fernanda C.G., 2018; Huang X K. R., 2024; Polderman TJ, 2015). Even though family data may have great value, collecting it remains a significant challenge. In research and clinical care, family data is often collected in a form of a ‘family history.’ Family histories can be structured or unstructured, collected through active questioning or via survey tools, and often focus on disease states for immediate family members. However, acquiring these family structures can be difficult. A typical family history can be labor intensive, is disease limited, can become outdated as soon as it is collected, and almost always relies on a person’s memory and awareness.

Recent research has demonstrated family data can be readily and reliably extracted from an electronic health record (EHR). These electronic pedigrees are called ‘e-pedigrees.’ Fernanda et al developed an e-pedigree algorithm using emergency contact information in an EHR. These have since been used to measure heritability for 500 disease phenotypes. (Fernanda C.G., 2018) Huang et al (Huang X E. R., 2018) demonstrated e-pedigrees can be generated using basic demographic data in an EHR (e.g., names and addresses) and can be applied to genetic mapping studies and the study of inheritance patterns. These two e-pedigree algorithms have since been unified (Huang X T. N., 2021).

Unfortunately, EHRs are temporally bounded thus limiting the depth of health and familial data available in e-pedigrees. More specifically, older generations, who inherently have richer health histories than younger generations, are often not linked to e-pedigree data. To address this, additional historical data is required.

In the United States, obituaries are often published as paid memorial announcements. The overall content of an obituary has remained constant for many decades. They often include name, date of birth, and date of death of the decedent. Obituaries may include information about memorial services and some individualized content (e.g., decedent’s job history). More importantly, obituaries regularly list decedent’s surviving and non-surviving (i.e., ‘preceding in death’) family members. This includes names and familial relationships.

Obituaries are good candidates for natural language processing (NLP) given their formulaic structure. There have been some attempts to extract family information from obituaries for biomedical research. He et al. developed a neural network model that extracted names and family relationships from 1,700 obituaries in the Minneapolis and Saint Paul, Minnesota area. (He K, 2021) Mumtaz and Qadar tried a rule-based system using regular expressions to extract family relations from text data. (Mumtaz, 2022) Rybinski et al. used obituaries to identify disease family history information. (Rybinski M, 2021) These studies stopped short and did not link family data to rich longitudinal health history data from an EHR. Other studies have used obituary data to find missing information in EHR patients, such as mortality status, but do not try to link to family data. (Ziedan, 2022)(El Emam, 2013).

We hypothesized family pedigrees obtained from obituary data can be successfully mapped to patients in an EHR. To test this hypothesis, we evaluated nearly 570,000 decedents with millions of family members extracted from obituaries published in the state of Wisconsin and cross referenced with Marshfield Clinic Health System’s (MCHS) EHR. In doing so, older, and more complete e-pedigrees can be linked to longitudinal health data.

## Methods

### Obituary Data

Access to obituaries were purchased from NewsBank Inc. (Newbank, 2023) Newsbank Inc. is a service that consolidates current and archived sources such as newspapers, newswires, business journals, periodicals, government documents and others. In this study, we limited our scope to publications in the state of Wisconsin. There were 1,360,088 articles from 81 sources (80 newspapers and one television station website). The date ranges were from March 19, 1989 through June 28, 2022. Not all provided articles were true obituaries. Some articles included simple death notices or news stories reporting on a death (e.g., automobile accidents); these were discarded as the NLP extract from them were erratic or provided no family information. Each obituary was provided in a separate file in XML format. On occasion, the same obituary for the same decedent was published in multiple sources. Only one was considered. In total, there were 567,279 obituaries for analysis.

### Natural Language Processing (NLP)

A Python program (version 3.6) was written to extract information of interest from the obituaries. The nltk package was used for basic NLP functions, such as word tokenizing. Given that the files were in XML, certain pieces of information could be found between XML tags in the file. These were the news source, date of publication, and the decedents full name. The rest of the information was in the main body of the text, and this had to be extracted using NLP methods to search for relationship keywords or phrases of interest (e.g., “survived by”). The program relied on common patterns that appeared in most obituaries. The first sentence often gave the name of the decedent, date of death (DOD), date of birth (DOB), and the decedents parents. Key phrases such as “died on”, “born”, and “born to” were searched for to identify information of interest. The Python package dateutil was used to identify dates. If a year was not given, such as “died on Wednesday, December 28”, date of publication was used to infer year of death. For the hypothetical example given above, if the obituary was published in January 2023, it was assumed the decedent died on Wednesday, December 28^th^, 2022. In cases where birth or death date could not be identified, usually due to poetic wording by the author, the minimum and maximum dates in the obituary were pulled and used for the missing fields. This worked well for birth date, as it was almost always the earliest date in the obituary. For the date of death, it could be off by a few days as it could pick up the date of the memorial service. This was mitigated by removing text that came after a phrase such as “services for…”, since these were usually at the end of the obituary.

Other than parents and possible spouses, most of the relatives were found at the end of the obituary, usually after the phrases “survived by,” or “preceded in death.” To identify these, logic was developed by looking for a relationship type in a predefined list of relationships, and then the names that followed the relationship type. Names were identified using the Stanford Named Entity Recognizer (NER), which classifies words into categories: person, location, organization, or O (other). The algorithm would identify a relationship type, and then extract names until it hit another known relationship type or the end of the sentence. Further procedures were applied to get first and last names along with possible spouses. From this, family data was extracted including relative’s name, type of relationship, and whether they were living at the time of the obituary as inferred by the “survived by” or “preceded in death” phrases.

### Linking Obituaries to EHR

To link the decedent to a patient in MCHS’s EHR, first name, last name, DOB, and DOD were all considered using different combinations of matching criteria. To increase precision, matches were further reduced if the individual had health records after the date of death as reported in the obituary. If more than one match occurred between obituary and patient in the EHR, the matches were discarded. Heritability of adult height was quantified as described subsequently (Evaluating accuracy of obituary pedigrees section) to evaluate decedent matching effectiveness. It was expected heritability would be lower if a matching strategy for decedents introduced error. There were no significant differences in heritability estimates between any of the five approaches, but there were large differences in number of matches (**Table 1**). As such, the approach that allowed the greatest number of decedent-patient matches (i.e., matched by last name and any two of first name, DOB, and/or DOD) was used for all follow-up analyses.

**Table 1.**
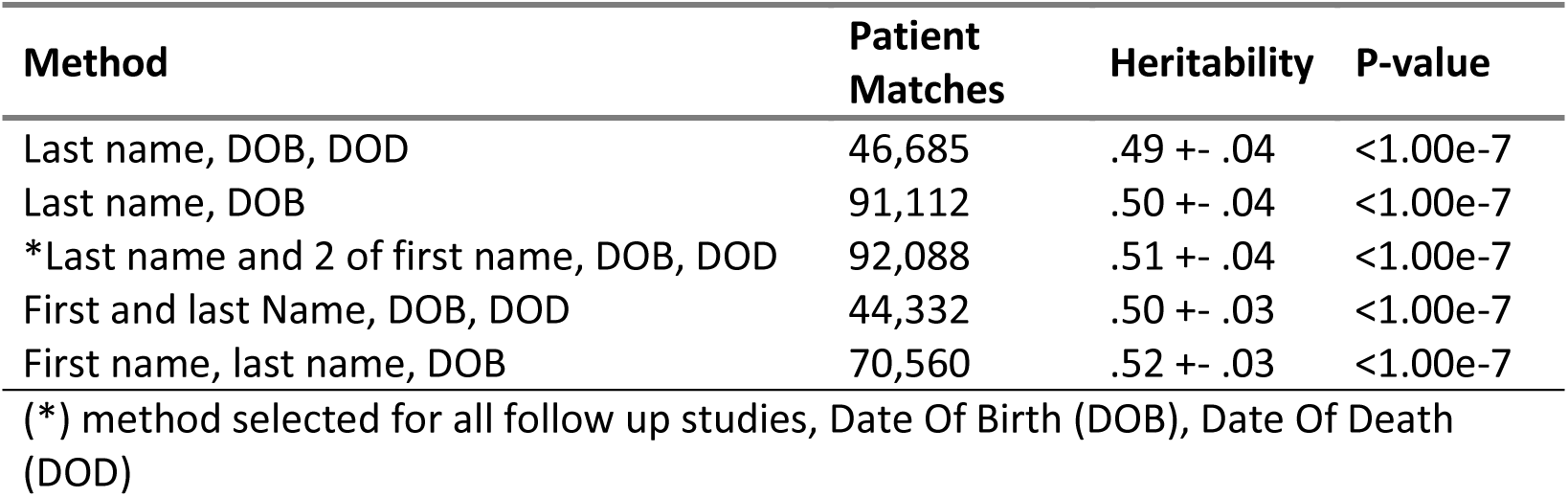
Heritability for the different methods for matching the obituary decedent to an MCHS patient.

For family members, only name, relationship, and mortality status were available. Names were matched to patients in the EHR, and then checked against their mortality status. Although date of birth was not available for family members, familial relationship to decedent was. This allowed us to approximate year of birth for family members. We assumed parents were 18-45 years older, children were 18-45 years younger, grandchildren were 30-55 years younger, and siblings and spouses were within 20 years relative to the decedent. As with the decedents, multiple matches were discarded. This was more common given limited matching criteria.

### Evaluating accuracy of obituary-pedigrees

Different methods and sources were used to evaluate obituary-pedigrees. Genomic data collected from participants who enrolled in the Personalized Medicine Research Project (PMRP) was used to define ground truth genetic relationships and compared to those that overlapped with obituary-pedigrees. PMRP consisted of 20,000 adult Marshfield Clinic patients that lived in a 19-zip code region surrounding Marshfield Wisconsin. The average age of participants was 63.9 and had on average 33 years of EHR data. Most PMRP participants have been genotyped on multiple whole-genome SNP arrays (Allaire P, 2023) and used to quantify a relatedness coefficient using VCF tools (Version 0.1.15) (Petr Danecek, 2011).

Obituary pedigrees were also compared to pre-existing e-pedigree data described previously (Huang X E. R., 2018) (Huang X T. N., 2021). At the time of these experiments, 579,561 MCHS patients could be placed into 173,368 e-pedigrees. Familial relationships between overlapping pedigrees were compared. Moreover, e-pedigrees were used to model error rates as inferred by heritability of adult height. In this instance 35,125 individuals from 14,870 families with available adult height measurements extracted from the EHR were randomly selected from the larger e-pedigree dataset. The number of families selected were based on input limitations of the software used to quantify heritability (i.e., S.A.G.E). To model error in the pedigree structures, we introduced randomness in the adult heights separately in males and females. For example, 5% of male heights were randomly permuted across all males and 5% of female heights were randomly permuted across all females. Coupled with the height included age of the person. This permutation introduced error in the pedigree relationships while maintaining the family structures and age and sex specific factors that influence adult height. This permutation was repeated 10 times for each percentage. Heritability of adult height was quantified using the ASSOC method in the S.A.G.E program. (S.A.G.E., 2021) Age and sex were included as covariates. Also included as a covariate was an age-by-age interaction term to account for the non-linear relationship between height and age as recommended by S.A.G.E. developer (see Acknowledgments).

## Results

### Pedigree structure

There were 567,279 obituary documents published in the state of Wisconsin available from NewsBank. Of these, decedents first name, last name, DOB, and DOD could be extracted from 86.5%, 86.5%, 79.2%, and 98.8% of the documents, respectively. Furthermore, 49.8% of the articles had family data that could be extracted totaling 8,166,534 family members. The most common relationships included parents, siblings, and children; 76%, 73%, and 70% of the obituaries had at least one person named, respectively.

MCHS serves predominantly patients living in central and northern Wisconsin. Newsbank had obituary data from 81 publishing groups throughout Wisconsin including those in large and small city markets. Of the 81 publishing groups, 18 papers were in a county where MCHS provides service. The largest number of patients that could be matched to obituary data came from these 18 papers, as expected. Likewise, the proportion of all obituaries evaluated, where the decedent could be mapped to a patient in the EHR, came from a paper published in MCHS service area (**Figure 1**). For example, the Milwaukee Journal Sentinel is likely the most read paper published in Milwaukee Wisconsin. Milwaukee is the largest city in the state but well outside MCHS service area. Of the 133,276 obituaries extracted from the Milwaukee Journal Sentinel between 1990-2022, 1% (n=1,199) had a decedent mapped to a patient in MCHS’s EHR. In comparison, Chippewa Falls, which is in the MCHS service area, had 22,733 extracted obituaries. Decedents that mapped to MCHS’s EHR made up 25% (n=5,683) of the obituaries.

**Figure 1.**
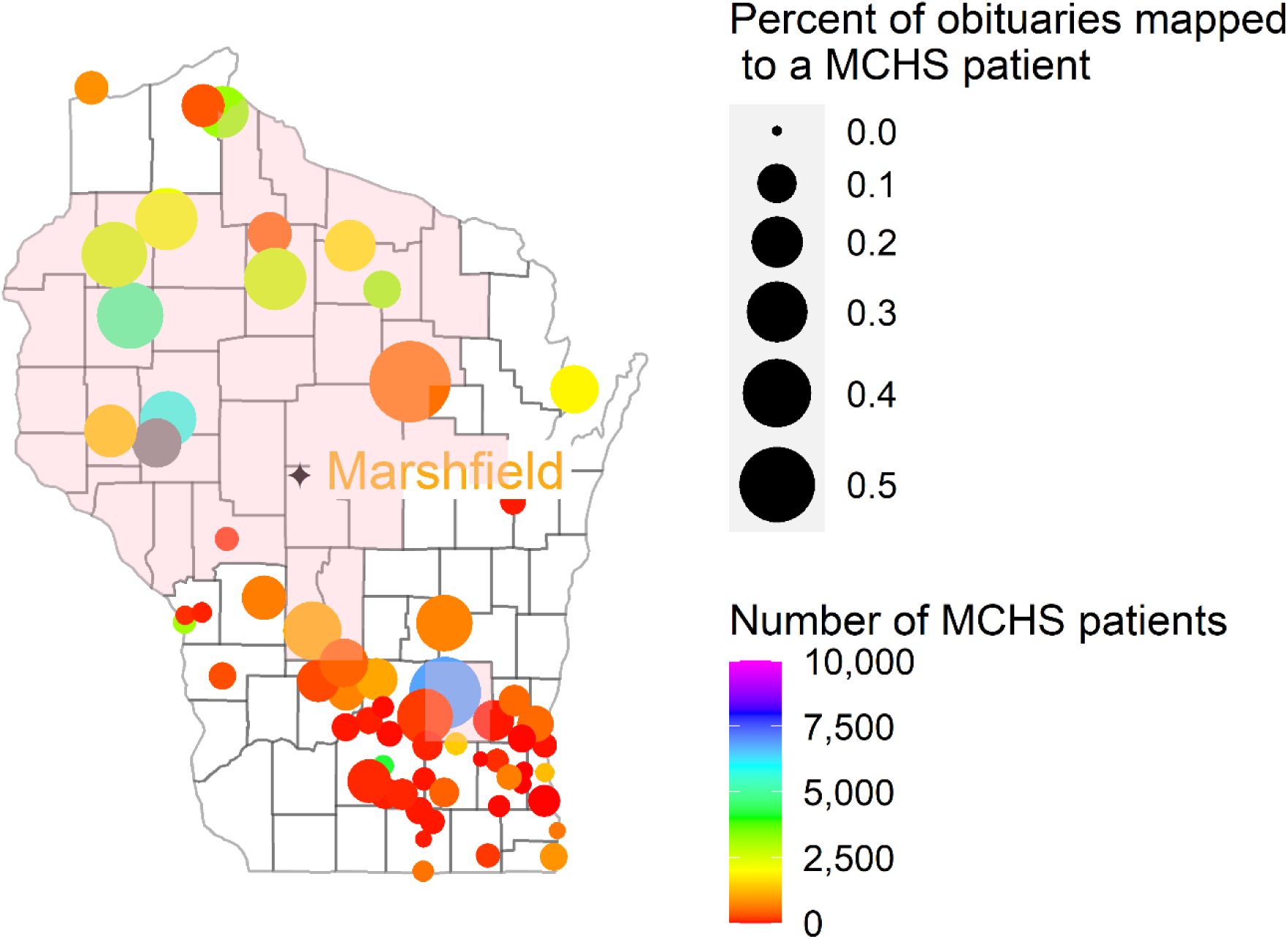
Map of Wisconsin containing locations of 81 publishers of obituaries. Shaded counties reflect MCHS service area. Included are number of decedents that are MCHS patients and proportion of all decedents linked to MCHS patients.

After matching decedent and family members to patients in the EHR, we identified 109,365 unique patients that were put into 34,158 pedigrees. Of all 44,332 decedents that could be matched to a patient in the EHR and were part of a pedigree, 18%, 48%, and 22% had a spouse, child, or grandchild that could also be linked to a patient in the EHR, respectively. This is compared to older generations where 14%, 4%, and 3% of the decedents had a mother, father, or grandparent linked to a patient in the EHR, respectively (**Figure 2a**). The largest pedigree consisted of twenty-one family members. Family sizes of six or more made up 11%. Whereas 50% of families were horizontal in structure consisting of only the decedent with a spouse and/or sibling(s), 50% had vertical structures consisting of two or more generations. **Figure 2b** shows the age at death distribution of the decedents.

**Figure 2.**
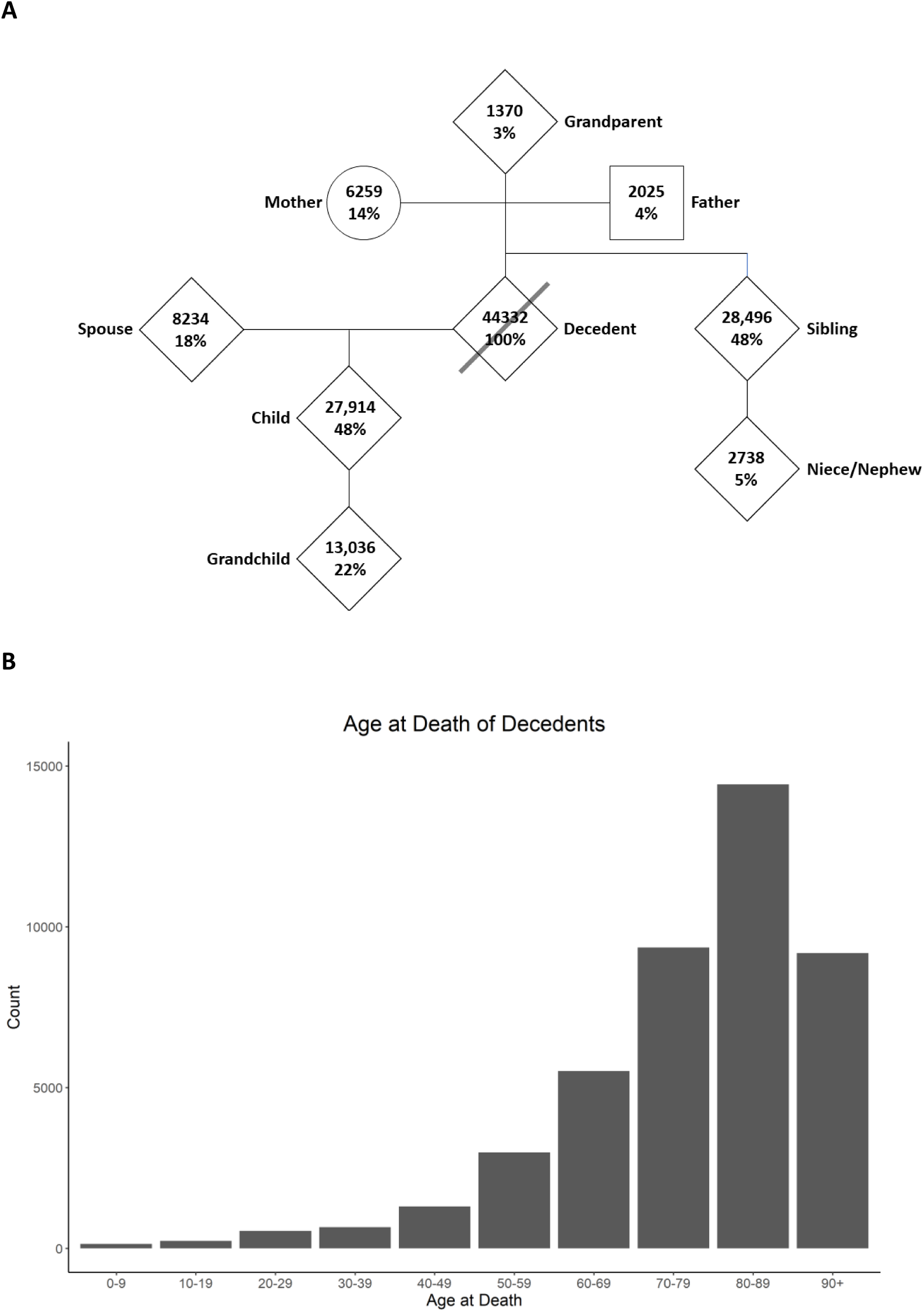
A) Summary of all 34,158 obituary-pedigrees linked to patients in MCHS. Circles are females, squares are males, and dimonds are unisex. Each symbol contains total number of patients in each relationship category and percentage of families that have that relationship defined. B) Distribution of the age of death of the decedents for whom a medical record was found in the EHR.

### Obituary-pedigree assessment

Currently, 579,561 MCHS patients can be placed into 173,368 highly accurate e-pedigrees (Huang X T. N., 2021). We compared obituary-derived pedigrees with e-pedigrees with focus on overlap. When two individuals were placed in a shared family in both datasets, relationship concordance relative to the decedent was 97.3%. **Table 2** shows the frequency with which the obituary-pedigrees and e-pedigrees were concordant by relationship type. Forty-six percent of concordant relationships consisted of children of the decedent (i.e., sons and daughters). This is expected given e-pedigrees are built from parent/child relationships. **Table 2** also shows discordances that were found. The most common was husbands, making up 24% of discordances. E-pedigrees classified these as either sibling or parent/child. Most of the other discordances were the result of generation mismatches (i.e., obituary data indicating a grandchild where e-pedigrees had the person as a child).

**Table 2.** Breakdown of relationships that were concordant and discordant between obituary and e-pedigree data.

| Relationship relative to decedent | Concordances |  | Discordances |  |
| --- | --- | --- | --- | --- |
|  | Count | Percent | Count | Percent |
| child | 1,104 | 76.2 | 1 | 2.44 |
| parent | 227 | 15.7 |  |  |
| sibling | 89 | 6.15 | 1 | 2.44 |
| grandchild | 28 | 1.93 | 9 | 22.0 |
| aunt/uncle |  |  | 1 | 2.44 |
| grandparent |  |  | 7 | 17.1 |
| nephew/niece |  |  | 2 | 4.88 |
| parent-in-law |  |  | 2 | 4.88 |
| spouse |  |  | 10 | 24.4 |
| stepchild |  |  | 8 | 19.5 |

In an effort to evaluate error in the obituary-defined pedigrees, we calculated a relatedness coefficient from pairs of individuals in the obituary dataset that also overlapped MCHS patients enrolled in PMRP (McCarty CA, 2005) where genomic data can provide ground truth (**Figure 3**). First-degree relationships should have a relatedness coefficient close to 0.25, whereas second-degree relationships (i.e., grandparents-grandchild, and aunt/uncle-niece/nephew) should be close to 0.125. Presumed unrelateds (i.e., spouses) would be close to zero. When comparing relatedness coefficients based on ground truth genomic data, 40 out of the 44 pairs were concordant. When assessing the four discrepancies, all were in first-degree relationships based on obituary data. One of the four pairs had a higher relatedness coefficient compared to what was expected. This pair were monozygotic twins. In other words, obituary-pedigrees correctly identified this pair’s social relationship (i.e., sibling) but not the genetic relationship. For the remaining three pairs, all had relatedness coefficients near 0 based on genetic data. Manual review of these three pairs did not provide further insights on cause of error. The totality of this result suggested genetically predicted relationships through obituary-pedigrees linked to an EHR had an accuracy rate of 90.9% or an error rate of 9.1%.

**Figure 3.**
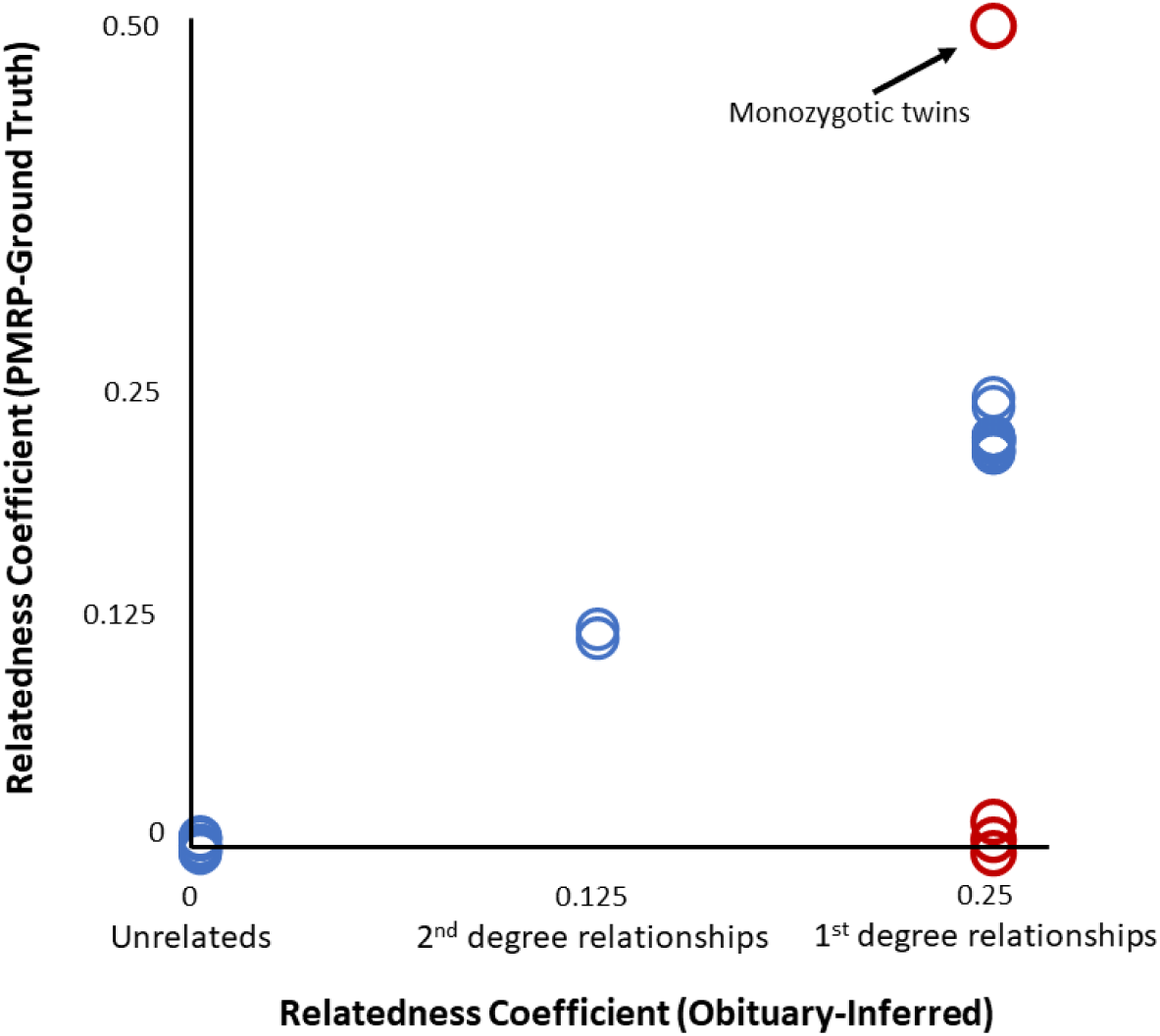
Comparison of relatedness coefficients from ground truth genomic data and inferred from obituary-pedigrees. Blue datapoints are concordant values. Red datapoints are errors from obituary-pedigrees.

Recognizing accuracy estimates from ground truth genomic data in PMRP came from a small sample of overlapping pairs of individuals, we devised a second strategy using existing e-pedigrees (Huang X E. R., 2018) (Huang X T. N., 2021) under the assumption there will be an inverse relationship between error in family relationships and heritability of adult height. In this instance, heritability of adult height in e-pedigrees was 0.70 +/- 0.01, consistent with other heritability estimates of this well characterized phenotype (Fernanda C.G., 2018). This is compared to 0.51 +/- 0.04 in obituary-pedigrees (**Table 1**). When artificially introducing random error in e-pedigree relationships, heritability decreased as predicted. Using this decay rate in heritability, we inferred a relative error rate in obituary-pedigrees of 15.3% [95%CI 11.7% – 18.9%] (**Figure 4**). This is close to the 9.1% rate observed from ground truth genomic data in PMRP.

**Figure 4.**
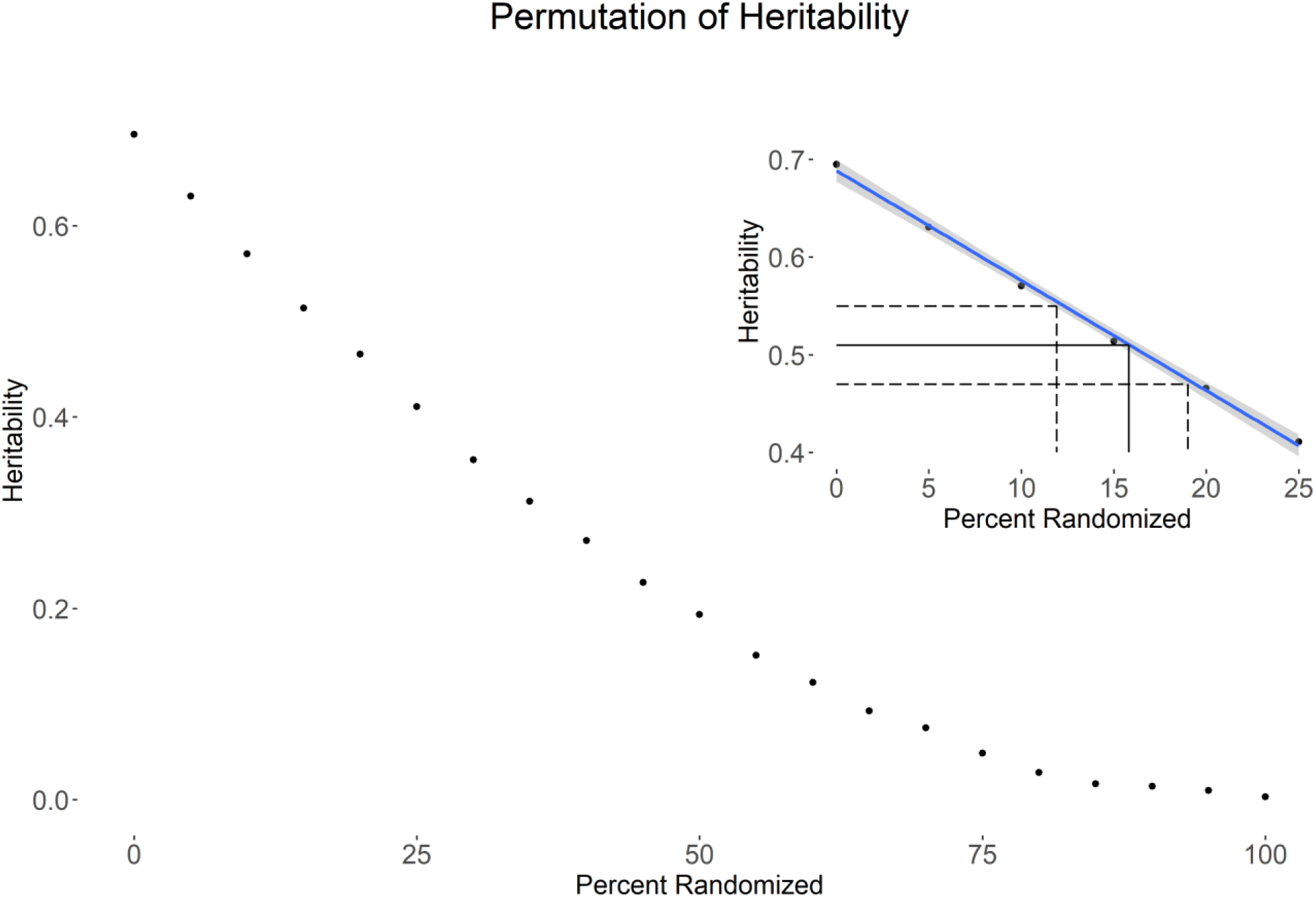
Results from the permutation experiment where heritability of adult height was measured in e-pedigrees under different conditions of randomness. The upper right is a zoomed in picture of the permutation experiment including a best fit line in the region of interest and 95% CI in gray. Overlayed is the observed heritability estimate calculated from obituary-pedigrees (H^2^=0.51 +- .04).

## Discussion

This study presents the first attempt to mine obituary data for family information and connect it to EHR data for use in research. This is especially valuable for older generations who cannot be readily linked to family data via e-pedigree algorithms (Huang X E. R., 2018) (Huang X T. N., 2021). Obituaries are a rich source of family pedigree data, and their consistent structure makes them a great source for standard NLP techniques resulting in 109,365 unique patients that were put in 34,158 pedigrees linked to longitudinal EHR data.

Two types of individuals extracted from an obituary need to be mapped to an EHR: decedent and their family members. We suspect mapping the decedent to an EHR is highly accurate given date of death and/or date of birth are explicit variables available from most obituaries. Moreover, using name and DOB is often all that is needed for patient identity verification in clinical care. Mapping family members to an EHR was far more difficult. This is because no discrete dates were available for family members. Common surnames may be underrepresented. The remaining unmatched family members either went by a different name in the EHR, were never patients of MCHS, or were not captured in the EHR due to its temporal nature. The latter likely occurred frequently since many of the family members who did not map to a patient in MCHS were parents of the decedent and likely predated the EHR.

Accuracy of our methods was difficult to ascertain given incomplete ground truth data. With the limited genomic data linked to obituary-pedigrees, 90.9% of genetic relationships were accurately predicted. This analysis also highlighted obituaries cannot discriminate monozygotic verses dizygotic twins. They will also struggle defining adopted relationships. When directly comparing familial relationships in e-pedigree verses obituary-pedigrees, a large percent of familial relationships (97.3%) were concordant. Given high agreements between these sources (**Table 2**), obituaries may be used to improve existing e-pedigrees. Lastly, we assessed accuracy by measuring heritability of adult height. Heritability of adult height in obituary-pedigrees was 0.51 (**Table 1**) compared to 0.70 in e-pedigrees. When modeling error using e-pedigree data, obituary-pedigrees may have a relative error of 15.3% [95%CI 11.8% – 18.9%]. When considering the totality of this data coupled with ground truth genomic data, obituary-pedigrees can reliably map patients in an EHR with an accuracy rate likely between 80%-90%.

The use of obituary-pedigrees linked to an EHR may have broad applications. In research, we have already demonstrated obituary-pedigrees can be used to measure heritability of adult height, albeit with a slightly elevated error rate. Adult height is just one of thousands of phenotypes that can be extracted from an EHR. This is particularly relevant since obituary-pedigrees capture the oldest generations who have lived long enough to experience many age-dependent diseases. This is compared to family data from standard e-pedigrees that are usually limited to young and healthy families and may explain why there is little overlap between the two pedigree types. In the end, a logical step would be to merge obituary-with e-pedigrees. This is possible with the most recent e-pedigree algorithm that allows users to incorporate external data when generating e-pedigrees (Huang X T. N., 2021). Merging obituary informed family data into e-pedigrees will provide larger families that capture older generations (**Figure 2b**) that often have rich health histories but with a small cost of reduced accuracy in the pedigree structures.

Expanded pedigree data linked to an EHR could support many other types of research. For example, family-based recruitment is extremely difficult. This is because some families are identified and recruited through pure serendipity, especially when an entire family reaches out to the research community given interest in their own unique health history. In other instances, enrollment of families can be painstakingly iterative. For example, one person is enrolled where a family history is collected to identify parents, children, and siblings. Based on this family history, additional family member may be approached for enrollment and the processes repeats itself until no more family members are available to participate. With pedigrees linked to an EHR, the most interesting families can be easily mined for the most interesting phenotypes then targeted as a whole. Even when multiple family members are not required for enrollment, but the collection of a family history data is still necessary, pedigrees connected to an EHR can allow researchers to capture some family history information without the dependency of a participant’s memory or awareness. With further improvements to this family in an EHR, it is conceivable these family histories may be used in clinical care allowing health systems to evaluate patient risk for many diseases.

This study had some limitations. First, the NLP method relied heavily on Stanford’s Named Entity Recognizer for identifying proper names. While this allows for identification of individuals, locations, organizations, etc., it does have the problem that rare or uncommon names are missed. This is unfortunate as these names would likely match correctly to unique names in the EHR. The opposite problem exists with common names where many individuals may share such a name. Second, we relied on data from Newsbank given their willingness to partner in research. Newsbank captured obituary data across Wisconsin but did not have data on many publishers in MCHS service area including cities of Marshfield and Wausau, two hub locations in the health system. Regardless, our methods are simple and likely applicable in any EHR system. Logical follow-up studies could consider leveraging large-language models to extract decedent and family data from obituaries. Regardless, obituary data linked to an EHR will always be dependent on the longitudinal nature of the EHR. Health systems with recently developed EHRs serving transient patient populations will not likely have extensive family data that can be pulled from obituaries, but this limitation is partially temporal as EHR systems mature.

In conclusion, this study builds on a growing body of knowledge demonstrating obituaries are a valuable source of family data. More importantly, we show for the first time this family data can be reliably linked to an EHR with marginal errors in the family relationships. By combining obituary data with an EHR, families with the oldest generations and richest phenotypic data can be combined with existing e-pedigrees for biomedical research and possibly clinical care.

## Acknowledgments

The authors would like to thank Robert Elston’s gracious support in S.A.G.E and guidance in heritability experiments. We would also like to thank our generous patients and participants. This work was partially supported by the generous donation from the Sturm Family Foundation and the following NIH awards: 1UL1TR002373 and 1R01GM130715.

